# Stochastic and Heritable Responses in Cell-Size Regulation and Survival to pH Stress in *C. glutamicum*

**DOI:** 10.64898/2026.10.28.762374

**Authors:** César Nieto, Zahra Vahdat, Luisa Blöbaum, Alexander Grünberger, Abhyudai Singh

## Abstract

Bacterial survival in fluctuating environments depends on the ability of individual cells to maintain physiological homeostasis under stress, yet how single cells recover from transient pH fluctuations remains poorly understood. We tracked the cell-size dynamics and survival of *Corynebacterium glutamicum* cells exposed to short, strong acid (pH 4, 20 min) or alkaline (pH 11, 15 min) pulses. Acid stress considerably increases the size added during the stress cycle, whereas alkaline stress increases it only modestly. Nevertheless, surviving cells maintain their division strategy and restore size homeostasis in the next cycle. Coordination between relatives is short-lived: sister cells have similar cell-cycle durations after stress, but durations become uncorrelated once the last division shared by two cells precedes the stress by more than about 1.5 generations. Survival of acid stress favors larger cells and is largely stochastic, with no detectable lineage heritability. In contrast, survival of alkaline stress favors smaller cells and is transiently heritable: co-survival probability remains high among relatives separated by up to two generations. These findings provide insight into pH survival in *C. glutamicum* and present statistical methods to study cell-size regulation and survival under transient, severe environmental fluctuations.

## Introduction

Microorganisms are constantly subjected to fast and strong environmental fluctuations [1]. Specifically, variations in pH can disrupt transmembrane proton gradients, impair enzyme kinetics, and alter nutrient solubility [2–4]. To survive these transient shocks, individual cells employ different biophysical and biochemical mechanisms, including general stress responses [5]. Depending on the underlying mechanism, these strategies can show traits of stochastic tolerance or heritable physiological adaptation [6, 7]. Single-cell studies have shown that survival of a severe stress is rarely a uniform, population-wide outcome: genetically identical cells in the same environment differ in whether, and how fast, they recover [8, 9]. For pH stress in particular, single-cell studies have documented cell-to-cell differences in the acid-stress response and in acid survival in *Escherichia coli* and other bacteria [10–14].

Recent work motivated by the classical Luria–Delbrück fluctuation assay [15] has revealed preexisting heterogeneity within isogenic cell populations that *primes* some bacteria to survive a future stress such as a lethal dose of antibiotics [16–19], as also reported for cancer cells exposed to targeted drugs [20] and induced UV tolerance in *E. coli* [21]. Where priming exists, the underlying heterogeneity is transiently heritable (i.e., it persists for several cell generations before reverting to a different state) and has been mechanistically linked to epigenetic processes and/or stochastic regulation within gene regulatory networks [22–24]. This raises the question of how much of the cell-to-cell variability in pH-stress survival is random, how much is tied to other variable physiological traits such as cell size, and how much reflects a transiently heritable state shared by related cells. A direct way to separate these contributions is to follow lineages and ask whether related cells respond alike [8, 25–27].

An example of an organism that must cope with such sudden environmental changes is *Corynebacterium glutamicum* [28]. This Gram-positive actinobacterium is an important model organism for the large-scale production of amino acids and other bulk chemicals [29]. These cells are moderately alkali-tolerant neutralophiles that typically maintain a narrow cytoplasmic pH of 7.5 ± 0.5 across external pH values of 6.0 to 9.0 [28, 30, 31]. When external pH drops below 5.5, the cells experience a sudden collapse of this internal equilibrium [32]. These cells lack the conventional amino acid–dependent decarboxylase systems (such as glutamate or arginine decarboxylases) that most bacteria use to consume excess protons during acid stress [4]. Instead, pH homeostasis relies on alternative physiological workarounds. Because acidic environments trigger the production of toxic reactive oxygen species (ROS) [33], a key mechanism for rapid recovery involves mycothiol [34]. Mycothiol acts as a cellular protectant, shielding the cysteine and methionine residues of proteins from ROS damage through reversible S-mycothiolation, which allows rapid protein regeneration once the stress is removed [34]. This mechanism operates alongside other ROS-management systems, such as catalase (KatA) and DNA-protection proteins [28]. Furthermore, when the electrochemical gradient increases during acid stress, *C. glutamicum* stabilizes its proton motive force through potassium uptake via channels such as CglK [35]. In contrast, defense against alkaline stress requires a fundamentally different approach, as ROS protection is unnecessary under high-pH conditions. Instead, survival and pH homeostasis rely heavily on cation/proton antiporters (Mrp-type Na^+^/H^+^ antiporters such as Mrp1) that actively drive protons into the cell against the gradient to keep internal equilibrium [36, 37].

Previous research on *C. glutamicum* found that survival of extreme pH pulses shows significant cell-to-cell variability [38]. Individual cells show distinct phenotypes: cells that resume growth rapidly, those that enter a lag phase before recovering late, and those that fail to recover entirely. Because biosynthetic and metabolic capacity scale with cell size [39], cell-to-cell differences in size are a natural candidate source of variability in these outcomes. To regulate their size, cells are traditionally described by their division strategy [40–42]: the *adder* (adding a constant size between divisions), the *sizer* (dividing at a critical cell size), or the *timer* (dividing after a fixed cell-cycle duration). These phenomenological strategies can be linked to specific mechanisms, revealing how cells prioritize the synthesis of division-related regulators over other cellular processes [43–45]. These mechanisms, and therefore the division strategies, can be altered in stressful or changing environments [46–49].

In this article, we study the cell size regulation under stress that under both acid and alkaline stress surviving cells keep the *adder* division strategy while transiently increasing the size added during the stress cycle, with a substantial increase under acid stress and a more modest one under alkaline stress; in both cases, the added size returns to control levels in the first cycle after the stress. The first division after a pH pulse shows a random delay. To characterize the randomness and heritability of this delay, we compare the durations of the cell cycle spanning the stress between pairs of related cells, using the normalized mean squared difference (NMSD) [26]. To estimate the level of relatedness, we use the lineage distance, defined as the time from the last division shared by two cells to the onset of stress [26, 27, 50, 51]. Under both stresses, the NMSD increases with lineage distance and saturates after about 1.5 generations, as it does under control conditions. We then apply the same lineage approach to survival, comparing the baseline survival probability with the conditional probability that a cell survives given that a relative survived. Survival of acid stress is largely stochastic, with no detectable lineage heritability, whereas survival of alkaline stress is correlated among relatives separated by up to two generations. This correlation suggests that survival of alkaline stress depends on a transiently heritable physiological state that related cells share before the stress. Together, these findings provide new insight into the strategies that *C. glutamicum* uses to cope with fluctuating environments.

## Results

### pH-stress responses at the single-cell level

To study how single cells respond to pH pulses, we tracked individual *C. glutamicum* cells growing in a microfluidic single-cell cultivation (MSCC) device [38, 52]. In this device, cells grow as small colonies inside monolayer cultivation chambers. At the same time, a continuous flow supplies fresh medium, so that the medium can be switched rapidly and reproducibly between control and stress conditions (Fig. 1A). Cells were imaged by phase-contrast microscopy every 10 minutes. From these microscopy images, cell contours were segmented and tracked over time using the deep-learning software DeLTA [53], followed by manual verification (see Methods). For each cell, we could therefore follow its growth, record the times of its divisions (to within the 10-min imaging interval), and reconstruct its position in the lineage tree of the colony. Data analysis pipelines are publicly available in our repository [54].

**Figure 1:**
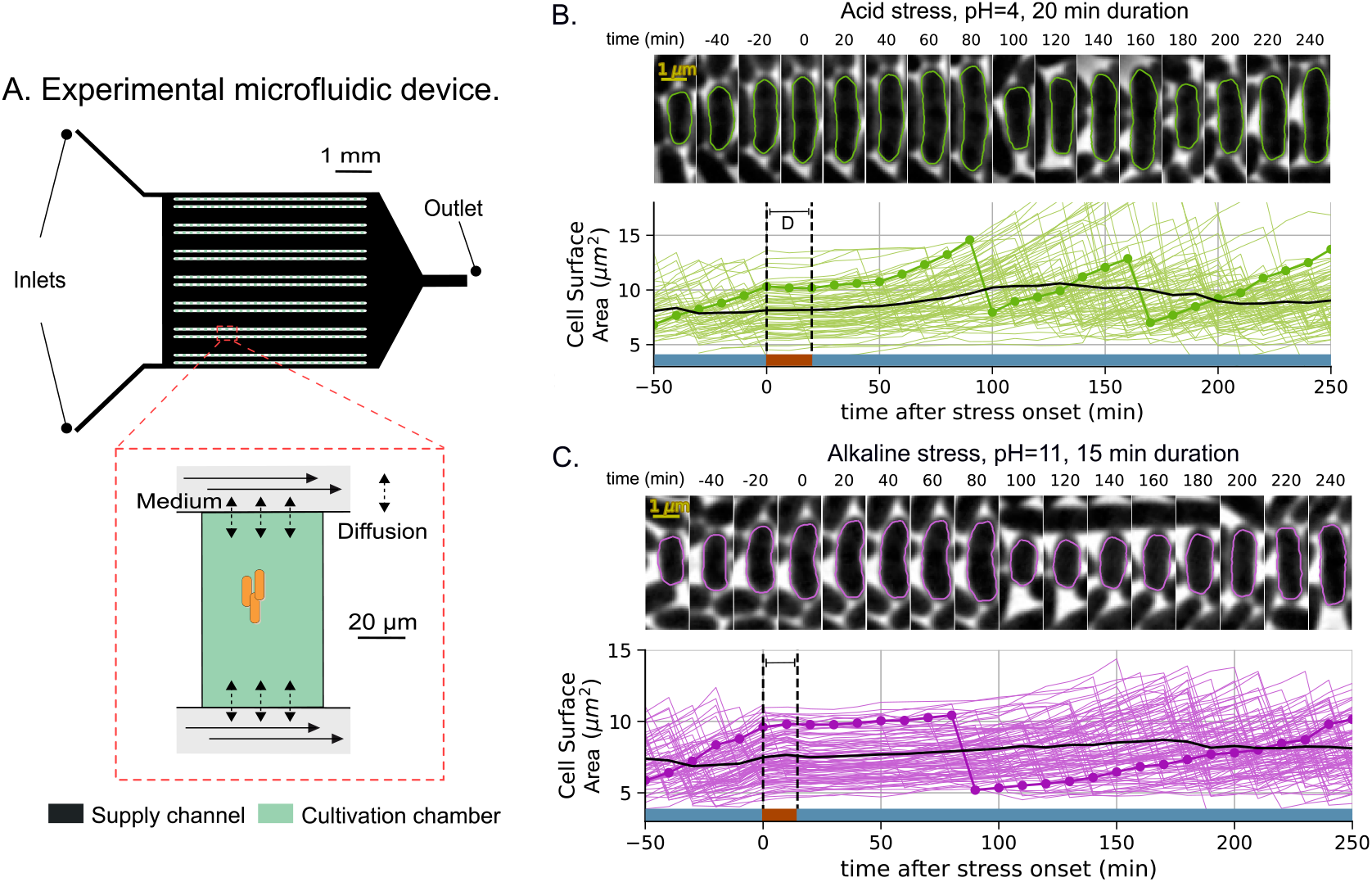
Experimental setup for studying the cell-size dynamics of *C. glutamicum* in response to pH stress. (**A**) The microfluidic single-cell cultivation (MSCC) device supplies a constant nutrient flow to 14 arrays of cultivation chambers using two inlets. (**B, C**) *Top*: examples of cell segmentation and tracking from microscopy images as a function of time. *Bottom*: cell-size trajectories over time for the studied cells. The cell shown on top is highlighted as a darker connected scatter plot. The black line represents the average cell size of all studied cells. The blue background marks time in control medium (pH 7); the red background marks the pH pulse. (**B**) Acid stress (pH 4, 20 min). (**C**) Alkaline stress (pH 11, 15 min).

We first characterized growth under control conditions in CGXII medium [55] at pH 7 (see Methods) and compared it with growth under two transient stress pulses: an acid pulse (pH 4, 20 min) and an alkaline pulse (pH 11, 15 min). As the readout of cell size, we used the cell surface area obtained from the measured cell projected area and cell length approximating cell shape to a capsule [56]. We chose surface area as cell size proxy because, in *C. glutamicum* and other cells, division is triggered by the accumulation of division precursors such as FtsZ with abundance proportional to cell surface area [57–60]. Therefore, throughout the paper, *cell size* refers to cell surface area.

To illustrate the cell-size dynamics, we show representative single-cell size trajectories (Fig. 1B,C, colored) together with the population average (solid black line). The blue background in Fig. 1B,C marks the periods of growth in control medium, and the red background marks the duration of the pH pulse. These trajectories reveal two contrasting responses. After acid stress, cells show a short delay before they resume growth, and some cells reach a relatively large size before dividing (Fig. 1B). After alkaline stress, cells instead show a long delay before growth resumes, and therefore a longer stress cycle (the cell cycle that spans the stress pulse), but grow to a cell size only slightly larger than before the stress (Fig. 1C). Next, we quantified these observations and asked how cell-size regulation, cell-cycle duration, and survival are affected by each pH pulse.

### *C. glutamicum* maintains an *adder* division strategy and recovers size homeostasis in the first post-stress cycle

Cell-size control is traditionally studied by estimating the division strategy, which describes how cells decide when to divide [40]. It can be quantified by the correlation between the newborn cell size *s*_*b*_ (the cell size at the beginning of the cell cycle) and the sizeΔ_*d*_ added during that cycle (Fig. 2D) [41, 42]. For the *adder* strategy, these two variables are uncorrelated. We verified that this is the case under control conditions (Fig. 2A). We then studied how the pH pulses perturb cell-size regulation and how cells recover in the following cycles.

**Figure 2:**
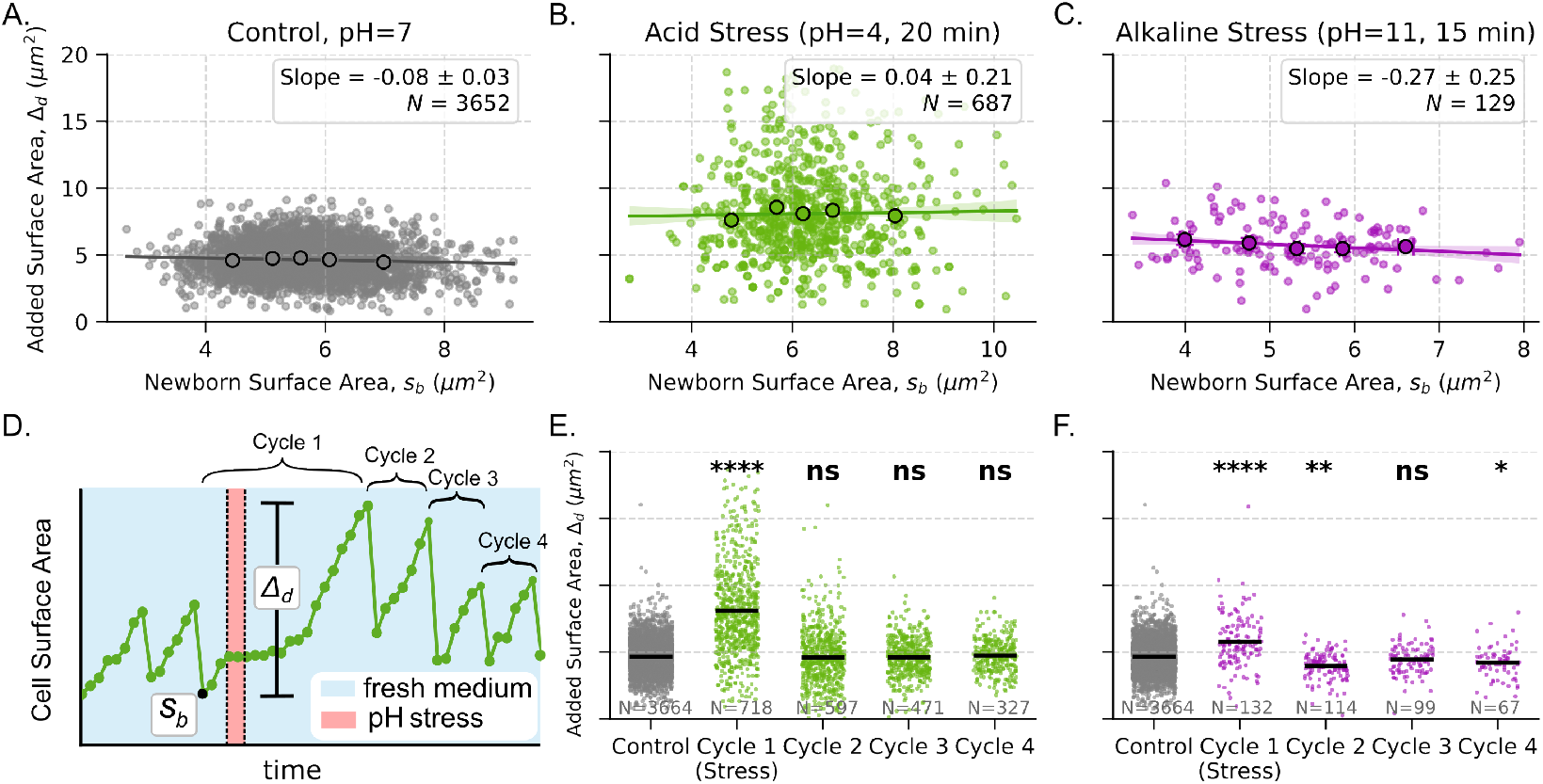
*C. glutamicum* maintains its *adder* division strategy through pH stress and restores size homeostasis in the first post-stress cycle. (**A**) Division strategy visualized as the relationship between the added sizeΔ_*d*_ during the cell cycle and the newborn cell size *s*_*b*_. These data correspond to control conditions (pH 7; 3652 points). (**B**) Division strategy for the stress cycle under acid stress (687 points). (**C**) Division strategy for the stress cycle under alkaline stress (129 points). (**D**) Cell-size dynamics of a cell under acid stress, showing the main variables studied. For the stress cycle, the newborn size *s*_*b*_ is the cell size at the beginning of the cycle that spans the pH pulse, and the added sizeΔ_*d*_ is the size added over that whole cycle, including the pulse. We also indicate the studied cycles: Cycle 1 is the stress cycle, and Cycles 2–4 are the post-stress cycles. (**E**) Jitter plot of the added size during and after acid stress, compared with control conditions (gray). The added size increases significantly during the stress cycle but recovers to control levels in the post-stress cycles. (**F**) Jitter plot of the added size during and after alkaline stress, compared with control conditions (gray). The added size shows a small but significant increase during the stress cycle and likewise recovers to control levels in the post-stress cycles. Statistical significance relative to the unperturbed control population was determined using Dunnett’s multiple-comparison test to control the family-wise error rate (^*\**^*p <* 0.05, ^*\*\**^*p <* 0.01, ^*\*\*\**^*p <* 0.001, ^*\*\*\*\**^*p <* 0.0001; ns, not significant) [61].

Under the *adder* strategy, cell-size control is summarized by the mean added size ⟨Δ_*d*_ ⟩, where the brackets ⟨·⟩ denote the average, and by its noise, quantified using the squared coefficient of variation 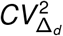, defined as the variance ofΔ_*d*_ divided by its squared mean ⟨Δ_*d*_⟩ ^2^. To organize the cell cycles relative to the pulse (Fig. 2D), we define the *stress cycle* (Cycle 1) as the cell cycle that spans the pH pulse, and the *post-stress cycles* (Cycles 2–4) as the ones that follow. For the stress cycle (Table 1), acid stress (pH 4, 20 min) produces a larger mean added size with greater noise than control conditions (Fig. 2B). In contrast, cells that survived alkaline stress show a modest increase in mean added size and a noise comparable to control conditions (Fig. 2C). Under both stresses, the added size of the stress cycle remains roughly uncorrelated with the newborn size, following the *adder* strategy (Fig. 2B,C). For the acid stress, the slopeΔ_*d*_ vs *s*_*b*_ is (0.04 ±0.21) while for the alkaline stress, it is (− 0.27 ±0.25). In the post-stress cycles, cells that survived either pulse rapidly restore normal size regulation: from Cycle 2 onward, the added size is indistinguishable from the control baseline (Fig. 2E,F), indicating robust and rapid homeostatic recovery.

**Table 1:** Mean added size ⟨Δ_*d*_ ⟩ and noise in added size 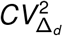 for the stress cycle (Cycle 1), compared with control conditions. Error intervals represent the 95% confidence interval.

| **Condition** | ⟨Δ<sub>*d*</sub>⟩ (*μ*m<sup>2</sup>) | *CV*<sup>2</sup><sub>Δ<sub>*d*</sub></sub> |
|---|---|---|
| Control (pH 7) | 4.65 ± 0.04 | 0.066 ± 0.003 |
| Acid stress (pH 4, 20 min) | 8.1 ± 0.2 | 0.18 ± 0.02 |
| Alkaline stress (pH 11, 15 min) | 5.7 ± 0.3 | 0.09 ± 0.03 |

### Lineage dependence of cell-cycle duration after stress

Our previous study gave a qualitative description of subpopulations with different, randomly variable recovery times, with little detail about the level of randomness and heritability [38]. Here, we quantify how the duration of the stress cycle depends on how closely two cells are related. We consider two cells to be related when they belong to the same colony (Fig. 3A), where a colony comprises all cells descending from a single cell present at the beginning of the experiment (the colony progenitor). After each division, one of the two daughter cells randomly keeps the label of the mother cell, and the other receives a new label. The relatedness of two cells is measured by their *lineage distance ℓ*: the time elapsed from the last division event shared by the two cells to the onset of the pH pulse (Fig. 3A). Lineage distance therefore measures how long before the stress two cells separated; sister cells have the shortest lineage distances. For instance, the two cells highlighted in Fig. 3A (orange and blue tracks) share a division event at time *t* ≈1.75 hours, even though each divides again afterward. Since the stress onset occurs at *t* = 4 hours, their lineage distance is *ℓ*≈2.25 hours. In control conditions as the cells maintain same conditions present before the stress, the lineage distance was measured relative to *t* = 4 hours. To obtain a general conceptual result, we also express lineage distance in generations, with one generation equal to the doubling time under control conditions (≈ 75 min).

**Figure 3:**
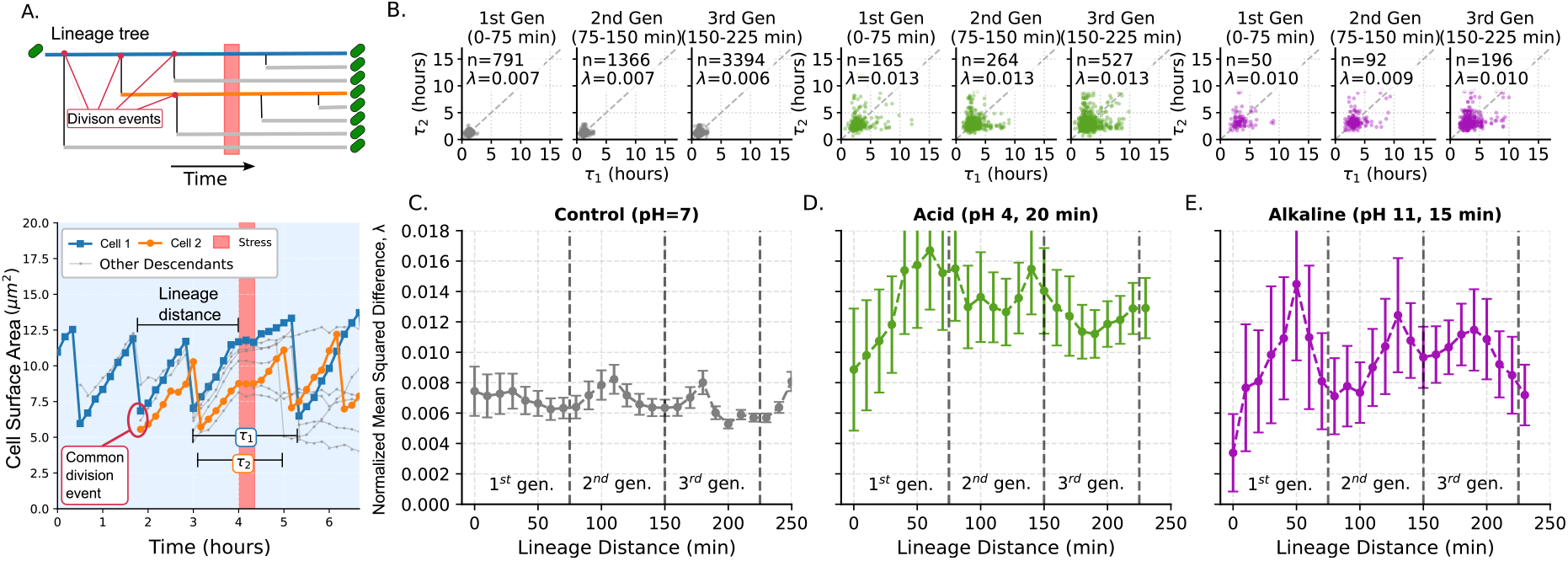
After stress, sister cells have similar cell-cycle durations, but this synchrony is progressively lost with lineage distance and vanishes beyond ≈1.5 generations (one generation ≈75 min). (**A**) Diagram of the studied variables. *Top*: an example lineage tree in which a selected cell (blue branch) has several relatives before stress (gray branches); a second cell that shares a division event with it is shown in orange. The lineage distance *ℓ*is defined as the time from the shared division event to the onset of stress (pink frame). Each branch point represents a division event. *Bottom*: cell-size trajectories of cells in the colony, with the cells from the top panel shown in their respective colors. We measured the duration of the stress cycle; *τ*_1_ and *τ*_2_ denote the cell-cycle durations of cells 1 and 2, respectively. (**B**) Examples of the relationship between *τ*_1_ and *τ*_2_ for three ranges of lineage distance: less than one generation (0–75 min), one to two generations (75–150 min), and two to three generations (150–225 min). The difference between the cell-cycle durations is measured by the normalized mean squared difference (NMSD) *λ* = ⟨ (*τ*_1_ − *τ*_2_)^2^⟩ */*(⟨ *τ*_1_⟩ ^2^ + ⟨*τ*_2_⟩ ^2^), with ⟨·⟩denoting the average. (**C–E**) NMSD as a function of lineage distance for (**C**) control conditions, (**D**) acid stress, and (**E**) alkaline stress.

To quantify the effect of pH stress on the timing of division, we measure the cell-cycle duration *τ* of the stress cycle. For any given pair of related cells, we denote the cell-cycle duration of the first cell *τ*_1_ and that of the second *τ*_2_. To make the comparison independent of which cell is labeled first, we symmetrized the data set, recording each pair twice: once as (*τ*_1_, *τ*_2_) and again as (*τ*_2_, *τ*_1_). This symmetrization guarantees that our estimates are invariant to cell ordering.

To measure how different *τ*_1_ and *τ*_2_ are, we use the mean squared difference of cell-cycle durations ⟨ (*τ*_1_ − *τ*_2_)^2^⟩. If two cells divide in synchrony, their cell-cycle durations are similar (*τ*_1_ ≈*τ*_2_); if they respond independently, the two durations can differ substantially. Because the mean cell-cycle duration differs between conditions, we normalize this quantity by the squared means, which gives the dimensionless normalized mean squared difference (NMSD, *λ*) between the paired durations [26, 27]:

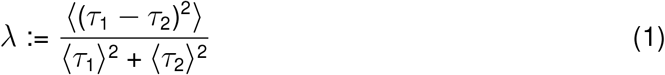

The NMSD is small when related cells divide in synchrony and grows as their cell-cycle durations diverge, so a larger *λ* indicates weaker synchrony. The NMSD depends not only on the correlation between the two durations but also on their overall spread, which itself changes under stress (Fig. 3B). This becomes explicit once we express *λ* in terms of the Pearson correlation coefficient between *τ*_1_ and *τ*_2_, denoted 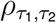, and the noise of the cell-cycle duration 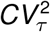 (see Methods):

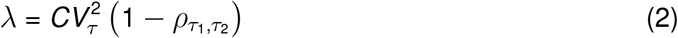

As a reference, we estimated the NMSD under control conditions (pH 7) showing small fluctuations around an asymptotic level (Fig. 3C). This behavior is also expected from simulations of exponentially growing cells dividing under the *adder* strategy (Fig. S1). In these simulations, the noise of the cell-cycle duration, 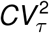, depends weakly on lineage distance, whereas the correlation 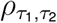 between the cell-cycle durations of related cells decreases from 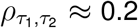 for the closest relatives to almost zero within about 1.5 generations. Accordingly, the NMSD is lowest for closely related cells (near-zero lineage distance), increases with lineage distance over approximately the first 1.5 generations, and then plateaus with possible oscillations related to the cell cycle effects (Fig. S1). The same trend is more pronounced for the stress cycle, for which *λ* plateaus at approximately twice the value observed under control conditions (Fig. 3D,E). These results support two main conclusions. First, the correlation between the cell-cycle durations of related cells vanishes for lineage distances beyond about 1.5 generations (Fig. S2), as in simulated cells under control conditions (Fig. S1). Second, stress cycles are not only longer (higher ⟨*τ*⟩) but also relatively more variable (higher 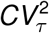).

We previously reported two classes of cell-cycle duration during recovery from pH stress: cells that recover relatively fast and cells that recover relatively slowly [38]. Here, our lineage analysis (Fig. 3B) shows that, under both stresses, the stress cycle length has less NMSD (*λ*) among cells with lineage distances below one generation. Cells with long stress cycles contribute substantially to the noise in cell-cycle duration resulting in a relatively high *λ*. They also make the estimate of *λ* fluctuate more between lineage-distance bins, as visible in Fig. 3D,E and Fig. S2. To obtain metrics robust to these outliers, we also estimated the Spearman correlation and a robust coefficient of variation (Fig. S3) [62]. The dependence of these metrics on lineage distance confirms that the correlation between the cell-cycle durations of related cells decreases with lineage distance, reaching zero beyond approximately 1.5 generations.

### Survival shows lineage heritability under alkaline stress but not under acid stress

As a final analysis, we investigated survival under the two stress conditions. Surviving cells in our data set are defined on the basis of cell-size dynamics after stress (Fig. 4A): cells that show no appreciable growth and no division during the rest of the time lapse (approximately 12 hours after stress) are considered non-surviving. Given the fate of each cell (surviving or non-surviving), we aimed to identify determinants of survival and to test whether survival is correlated between related cells. First, we compared the cell size at stress onset between surviving and non-surviving cells. The effect of cell size on survival depends on the stress: cells that survived acid stress were significantly larger on average, whereas cells that survived alkaline stress tended to be smaller than non-surviving ones (Fig. 4B).

**Figure 4:**
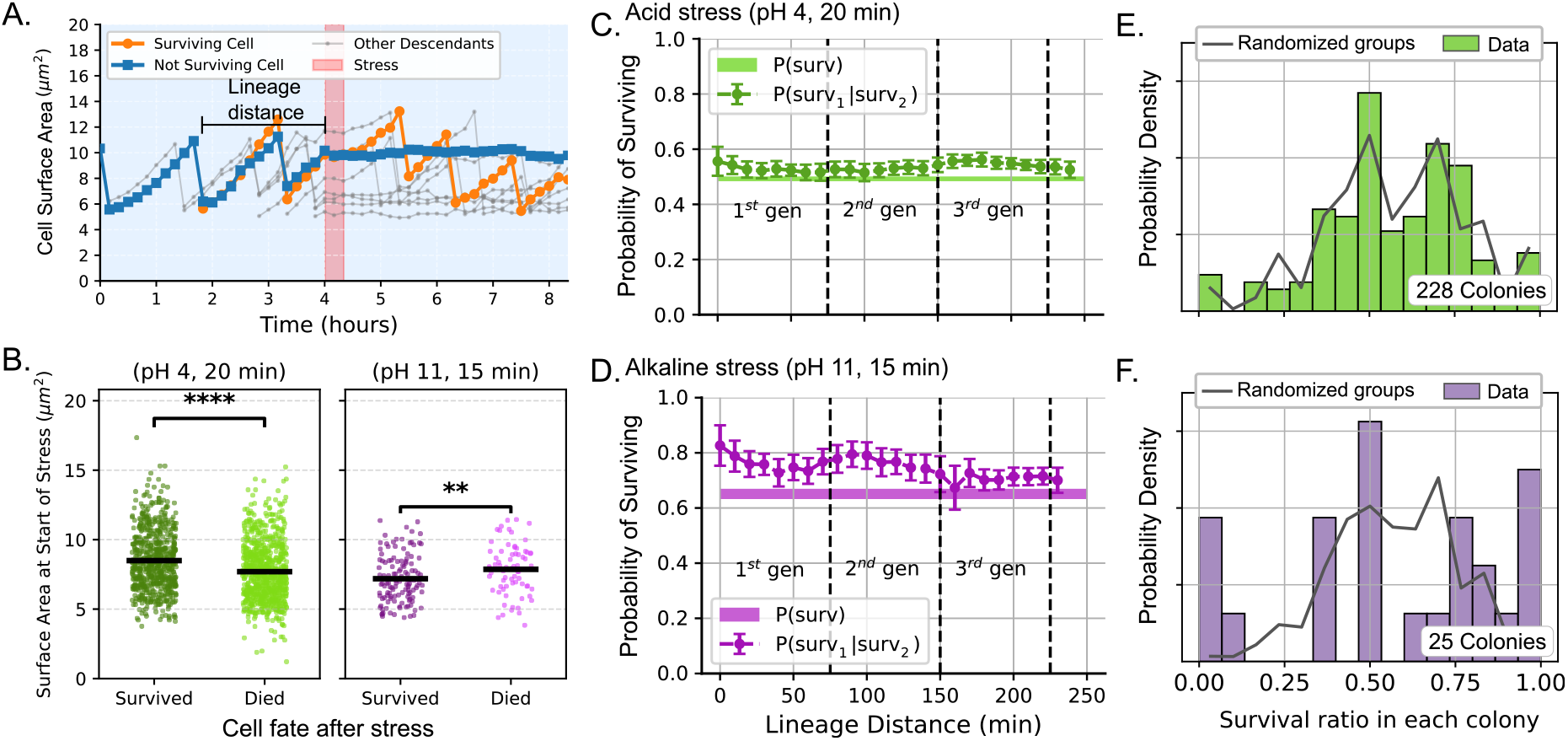
Survival of acid stress is largely stochastic and favors larger cells, whereas survival of alkaline stress favors smaller cells and is correlated among relatives separated by up to two generations. (**A**) Cell-size trajectories of a surviving cell (orange) and a non-surviving cell (blue) descended from the same cell. The lineage distance *ℓ*is measured from their last shared division to stress onset. Other cells of the colony are shown in the background (gray). (**B**) Dependence of survival on cell size at stress onset. Cells that survive acid stress are larger at stress onset, whereas cells that survive alkaline stress are smaller. Statistical significance was determined using Dunnett’s multiple-comparison test to control the family-wise error rate (^*\**^*p <* 0.05, ^*\*\**^*p <* 0.01, ^*\*\*\**^*p <* 0.001, ^*\*\*\*\**^*p <* 0.0001; ns, not significant) [61]. (**C**) Baseline survival probability *P*(surv) (horizontal line) and conditional survival probability *P*(surv_1_ | surv_2_) as a function of the lineage distance between the two cells, for acid stress. (**D**) The same analysis for alkaline stress. (**E**) Distribution of survival ratios across colonies after acid stress (histogram), compared with the distribution for randomized colonies (gray line). The acid-stress results are consistent with the absence of lineage heritability. (**F**) The same analysis for alkaline stress; the results support lineage heritability.

To test for lineage heritability of survival, we compared, as a function of lineage distance, the baseline survival probability, *P*(surv), which uses no information about relatives, with the conditional probability that a cell survives given that a related cell also survived, *P*(surv_1_ |surv_2_). These probabilities were estimated using the relative frequency of surviving cells (see Methods). For acid stress, *P*(surv_1_ |surv_2_) is almost indistinguishable from *P*(surv) at all lineage distances (Fig. 4C), suggesting that survival of acid stress is largely stochastic and lacks lineage heritability. Conversely, survival of alkaline stress shows lineage heritability: *P*(surv_1_ |surv_2_) remains higher (non-overlapping 95% confidence intervals) than *P*(surv) for relatives separated by less than two generations (Fig. 4D). Beyond two generations, *P*(surv_1_ |surv_2_) and *P*(surv) are practically equal.

A complementary way to quantify lineage heritability, in the spirit of the fluctuation test, is to analyze the distribution of survival ratios across colonies [17, 18, 63]. We define a colony’s survival ratio as the fraction of its cells that survive the stress. If survival is heritable, related cells share their fate and the survival ratio varies strongly from colony to colony. As a control with no heritability, we generated randomized colonies from the experimental data: for each colony, we drew as many cells as the colony contains from the pooled cells of all colonies, irrespective of lineage, and computed the survival ratio of this randomized colony (see Methods). Consistent with our analysis of survival probability versus lineage distance, the distribution of survival ratios under acid stress closely matches the randomized control (Fig. 4E). In contrast, as expected for a heritable state, the distribution of survival ratios under alkaline stress differs appreciably from that of the randomized colonies (Fig. 4F). Together, these results indicate that whereas survival of acid stress is largely stochastic, survival of alkaline stress in *C. glutamicum* is influenced by a transiently heritable physiological state shared among close relatives.

## Discussion

In this article, we studied the single-cell size dynamics and the survival of *C. glutamicum* cells in response to strong and short pH pulses. As observed through microfluidic single-cell cultivation combined with image-based cell tracking (Fig. 1), acid and alkaline stress trigger contrasting responses in cell size and in the lineage heritability of survival. Cells recovering from either pulse maintain an *adder* division strategy and restore the control added size in the first post-stress cycle (Fig. 2). In the stress cycle itself, acid stress considerably increases both the mean and the noise of the added size, whereas alkaline stress produces a modest increase in the mean with a noise comparable to control conditions.

The relatively large increase in both the mean and the noise of added size under acid stress probably results from a disruption of cell physiology that is absent under alkaline stress. We hypothesize that the larger added size under acid stress arises from a transient decoupling between growth and the cell-division machinery. Environmental stress can perturb the assembly of FtsZ filaments and delay division even as growth continues [64]. Because *C. glutamicum* lacks the amino acid decarboxylase systems that other bacteria use to survive extreme acid, the severe drop in internal pH likely forces a reallocation of energetic resources toward immediate survival [65]. Stabilizing the proton motive force through potassium uptake may then prolong this division delay while cell size continues to increase. The increased noise in added size is consistent with the stochastic nature of acid-induced damage: acidic environments trigger ROS production [33, 34], and both the oxidative-stress response and single-cell survival under stress vary strongly from cell to cell [11, 66, 67].

In contrast, mitigation of alkaline stress relies on dedicated effector proteins, specifically Mrp-type Na^+^/H^+^ antiporters [37], yielding a tighter, less variable recovery of the added size. Unlike acid-induced ROS damage, whose repair is multi-layered and highly variable, these membrane transporters manage alkaline stress efficiently without generating widespread oxidative damage [28]. Because expression states, such as an elevated level of membrane transporters, are passed from mother to daughter cells and are reset only over successive generations [68, 69], survival of an alkaline pulse can reflect a transiently heritable physiological state shared among close relatives. This difference between the two responses suggests that, when damage is diffuse and its repair depends on several general-purpose systems (as for acid stress, which engages mycothiol, catalase, DNA-protection proteins and potassium uptake) rather than on a single dedicated effector, survival behaves as a largely random variable with high cell-to-cell variability [70].

Our analysis of cell-cycle duration shows that the synchrony between related cells is lost once their last shared division precedes the stress by more than approximately 1.5 generations (Fig. 3). The same lineage dependence is observed and predicted by simulations of cells in control conditions, which indicates that the delay induced by a pH pulse is shared only between close relatives, to the same extent as cell-cycle duration in unstressed cells. Because lineage distance is measured before the pulse, this result describes how long a pre-stress physiological state remains shared between relatives, not how long the effect of the stress persists afterward. We propose that the determinants of cell-cycle duration are reset rapidly. Under the *adder* strategy, a deviation of the newborn size from the population average is halved at each generation [71], so the size that two relatives inherited from their common ancestor contributes little to their cell-cycle durations after one to two generations. In parallel, division regulators and other factors inherited from the common ancestor are diluted by growth: within 1.5 generations, cell size has more than doubled. By comparison, the state associated with survival of alkaline stress remains shared for somewhat longer, up to two generations (Fig. 4D).

Survival patterns differ clearly between the two stress conditions (Fig. 4). Survival of acid stress is largely stochastic, with a bias toward cells that are larger at stress onset and no detectable lineage heritability. A possible explanation for the size bias is that larger cells contain a larger pool of cytoplasmic scavengers, chaperones, and intact proteins, increasing the like-lihood that a cell survives and repairs oxidative damage [38, 72, 73]. Conversely, survival of alkaline stress favors cells that are smaller at stress onset and is associated with a transiently heritable physiological state that links the fates of relatives separated by up to two generations. In light of recent studies, the size bias may reflect the fact that smaller cells, having less cytoplasm to re-equilibrate, require less absolute energy and ion flux to restore internal pH homeostasis via Mrp-type Na^+^/H^+^ antiporters [74–76]. Furthermore, smaller cells, being far from their next division, may be able to prioritize the energetically demanding activity of these pumps over growth, making them more efficient at maintaining homeostasis under alkaline stress.

In summary, this article applies several statistical methods to analyze the response of *C. glutamicum* to strong pH pulses. Our results reveal that survival of acid stress is largely stochastic, whereas survival of alkaline stress shows lineage heritability. Under acid stress, larger cells have a higher survival probability, whereas under alkaline stress the surviving cells tend to be smaller. The duration of the stress cycle shows the same lineage dependence as cell-cycle duration under control conditions, albeit with greater noise. These results illustrate how individual bacteria combine immediate biophysical constraints with physiological states inherited from their recent ancestors to survive rapidly changing environments.

## Methods

### Biological preparation and nutrient medium

In this study, the bacterial strain *Corynebacterium glutamicum* ATCC 13032 was used [77]. *C. glutamicum* was cultivated in CGXII medium [78]. CGXII medium consists of the following components per liter of demineralized water: 20 g of (NH_4_)_2_SO_4_, 1 g of K_2_HPO_4_, 1 g of KH_2_PO_4_, 5 g of urea, 13.25 mg of CaCl_2_ ·2H_2_O, 0.25 g of MgSO_4_·7H_2_O, 10 mg of FeSO_4_·7H_2_O, 10 mg of MnSO_4_·H_2_O, 0.02 mg of NiCl_2_·6H_2_O, 0.313 mg of CuSO_4_·5H_2_O, 1 mg of ZnSO_4_·7H_2_O, 0.2 mg of biotin, and 40 g of D-glucose. The concentration of protocatechuic acid (PCA) in the medium was 30 mg/L. For shake-flask cultivation, 42 g/L MOPS buffer was added. The solutions were autoclaved or sterile-filtered. The medium was adjusted to the given pH using either H_3_PO_4_ (acid pulse) or NaOH (alkaline pulse). The adjusted medium was sterile-filtered again to remove contamination and pH-induced precipitates. All chemicals were purchased from Carl Roth.

For the preculture of *C. glutamicum*, 10 mL of CGXII medium at pH 7 in a 100-mL baffled shake flask was inoculated from a cryo-stock and cultivated overnight at 30 *°*C on a rotary shaker at 120 rpm. The main culture was prepared from this preculture, inoculated at an initial optical density (OD_600_) of ≈ 0.05. After reaching OD_600_ ≈ 0.2, the culture was transferred to a 1-mL syringe and the microfluidic chip was inoculated.

### Chip design and fabrication

For our microfluidic experiments, a microfluidic chip was adapted from previous work [79]. The chip consists of 14 arrays of cultivation chambers (80 *µ*m × 90 *µ*m × 650 nm) arranged in seven blocks. The supply channels have a height of 10 *µ*m and a width of 100 *µ*m (see Fig. 1A). The microfluidic chips were fabricated using soft lithography. Polydimethylsiloxane (PDMS) [80] was deposited on a silicon wafer at a 10:1 ratio of curing agent to linker (Sylgard 184 Silicone Elastomer, Dow Corning Corporation, USA). After degassing, the PDMS was baked at 80 ^*°*^C for 2 h. The PDMS chips were then cut from the wafer, and the inlets and outlets were punched (Reusable Biopsy Punch, 0.75 mm, WPI, USA). The PDMS chip and a glass slide (D 263 T eco, 39.5 ×34.5 ×0.175 mm, Schott, Germany) were cleaned three times with isopropanol, activated by an O_2_ plasma cleaner (Femto Plasma Cleaner, Diener Electronics, Ebhausen, Germany), and assembled. In a randomized process, the cultivation chambers were loaded with 1–5 cells. Once sufficient chambers were loaded, the flow of cell suspension was stopped and medium flow was started using high-precision pressure pumps (Line-up series, Fluigent, Jena, Germany). Pressures of 180 and 20 mbar were set to initiate medium flow. The detailed flow profile was published previously [38, 81].

### Live-cell imaging

Live-cell imaging was performed using an automated inverted microscope (Nikon Eclipse Ti2, Nikon, Germany). The microscope was placed in a cage incubator for optimal temperature control at 30 °C (Cage Incubator, OKO Touch, Okolab S.R.L., Italy). The microfluidic chip was attached to a custom chip holder. A 100× oil objective (CFI P-Apo DM Lambda 100 ×Oil, Nikon GmbH, Germany), a DS-Qi2 camera (Nikon GmbH, Germany), and an automatic focusing system (Nikon PFS, Nikon GmbH, Germany) were used to prevent thermal drift during cultivation. For each experiment, 150 cultivation chambers were manually selected using NIS Elements software (Nikon NIS Elements AR software package, Nikon GmbH, Germany). Phase-contrast images of each position were acquired every 10 min with an exposure time of 50 ms and a lamp intensity of 10%.

### Estimation of lineage distance

To quantify the relatedness of two cells within a colony, we defined the lineage distance *ℓ*as the time elapsed (in minutes) from the last division event shared by the two cells to the onset of stress. Lineages were reconstructed for each position and colony by computationally tracking mother–daughter relationships across successive frames. For any given pair of related cells within a colony, the lineage distance was obtained by tracing the branches of the lineage tree back to their last shared division event and measuring the time from that event to stress onset. For the control experiment, to maintain consistency, we estimate the lineage distance relative to the time *t* = 4h since the beginning of the experiment. For the NMSD analysis (Fig. 3C–E), cell pairs were aggregated into overlapping bins of ± 20 min. For obtaining the scatter plots, of Fig. 3B, pairs were grouped into ranges of one generation (approx 75 min).

### Normalized mean squared difference and its relationship to cycle correlation and variability

To quantify the difference between the cell-cycle durations of pairs of related cells, we defined the NMSD *λ* as in Eq. (1). The numerator ⟨ (*τ*_1_ − *τ*_2_)^2^ ⟩ is the mean squared difference between the cell-cycle durations (*τ*_1_ and *τ*_2_) of a pair of related cells, which expands as

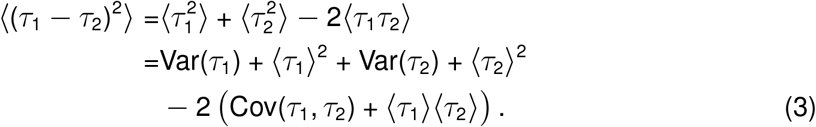

Because the data set is symmetrized by recording each pair in both orders, the two variables have identical statistics, giving equal means (⟨ *τ*_1_⟩ = ⟨ *τ*_2_⟩ = ⟨ *τ*⟩) and variances (Var(*τ*_1_) = Var(*τ*_2_) = Var(*τ*)), where *τ* is the cell-cycle duration without regard to ordering. Then

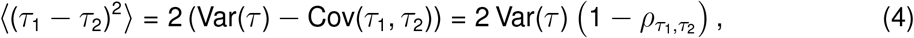

where 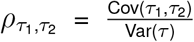 is the Pearson correlation coefficient between the paired cell-cycle durations, while the denominator collapses to ⟨*τ*_1_⟩^2^ + ⟨*τ*_2_⟩^2^ = 2⟨*τ* ⟩^2^. This simplifies the NMSD to

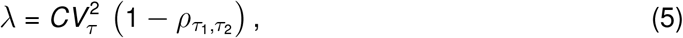

where 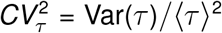 is the noise (squared coefficient of variation) of the cell-cycle duration.

### Survival probability and bootstrap estimation

To assess both the baseline survival and its correlation among related cells, we calculated the baseline (marginal) survival probability, *P*(surv), and the conditional survival probability, *P*(surv_1_ | surv_2_). To smooth the estimates across lineage distances, data points were aggregated using an overlapping moving window. For each evaluation distance *ℓ*(in steps of the inter-frame time of 10 minutes), cell pairs were pooled if their lineage distance fell within a window of ±30 minutes centered on *ℓ*. The window was truncated at 0 and at the maximum observed lineage distance. This moving-window approach allowed robust bootstrap estimation even in intervals with sparse pair counts. For each window, binary survival outcomes (1 for surviving, 0 for non-surviving) were extracted for all pairs. A bootstrap approach with 500 iterations was used to estimate the means and assess statistical variation. In each iteration, cell pairs were sampled with replacement to create a bootstrapped cohort. The baseline probability *P*(surv) was defined as the fraction of all surviving cells with no discrimination by lineage distance. The conditional probability *P*(surv_1_ | surv_2_) was computed as the number of pairs in which both cells survived, divided by the total number of surviving reference cells in that sample. The reported probabilities correspond to the mean over bootstrap iterations. Error bars represent the 95% confidence interval for the estimated probabilities [82].

### Distribution of survival ratios across colonies

To quantify lineage heritability of survival, we established a null model representing the distribution of survival ratios under the assumption of no lineage heritability (i.e., survival independent of ancestry). We generated this randomized control from our experimental data by resampling. First, we pooled all cells across all analyzed colonies. Next, for each analyzed colony, we counted its number of cells and randomly selected that same number from the pool, with replacement, regardless of colony of origin. We then used the fates of these randomly selected cells to compute the survival ratio of the randomized colony. This randomization was repeated 1000 times for the entire data set. The aggregated survival ratios of the randomized colonies formed the null distribution (shown as the randomized control in Fig. 4E and Fig. 4F). The empirical distribution of colony survival ratios was then plotted against this null distribution. A clear divergence of the empirical distribution from the randomized control indicates lineage heritability within colonies.

## Author contributions

- **Conceptualization:** A.G., A.S., C.N.
- **Investigation:** L.B.
- **Data Curation:** C.N., Z.V.
- **Formal Analysis:** C.N., Z.V., A.S.
- **Funding Acquisition:** A.S., A.G.
- **Methodology:** A.S., A.G.
- **Resources:** A.S., A.G.
- **Software:** C.N., Z.V.
- **Supervision:** A.S., A.G.
- **Visualization:** C.N., Z.V. and L.B.
- **Writing – Original Draft:** C.N., A.S.
- **Writing – Review & Editing:** A.S., L.B., A.G.

## Acknowledgments

L.B. and A.G. are supported by the Joachim-Herz-Foundation (Add-on Fellowship for Interdisciplinary Life Sciences). A.S. and C.N. acknowledge support from NIH-NIGMS through grant R35GM148351.

## Data Availability Statement

Experimental data and scripts for analysis are publicly available in a Zenodo repository: https://doi.org/10.5281/zenodo.23274001.

