## Supplementray Information for "Stochastic and Heritable Responses in Cell-Size Regulation and Survival to pH Stress in *C. glutamicum*"

### Supplementary Figures

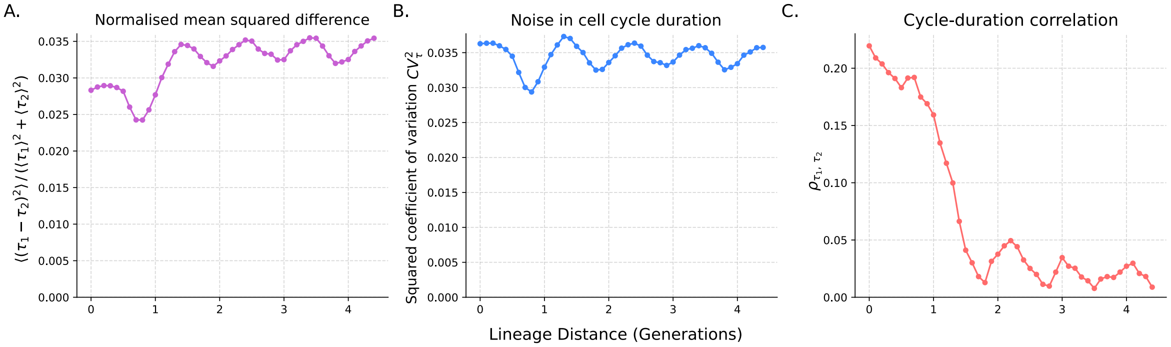


Figure S1: **Simulated decomposition of the normalized mean squared difference of cell-cycle durations into cell-cycle noise and cell-cycle correlation for steady-state cell populations.** The temporal evolution of cell-cycle dynamics is shown for simulated microcolonies undergoing exponential growth and dividing under an *adder* strategy. **(A)** Normalized mean squared difference ($\lambda$) of cell-cycle durations between lineage-related cell pairs as a function of lineage distance. **(B)** Baseline phenotypic noise in cell-cycle duration, quantified by the squared coefficient of variation ($CV^2_\tau$). **(C)** Pearson correlation coefficient ($\rho_{\tau_1,\tau_2}$) of cycle durations between related cells, illustrating the decay of transgenerational memory in a control steady-state system. Data points represent 500 colonies evolving from a single cell over 4.5 generations. Time statistics were estimated using a moving average with a $\pm$0.25-generation sliding window.

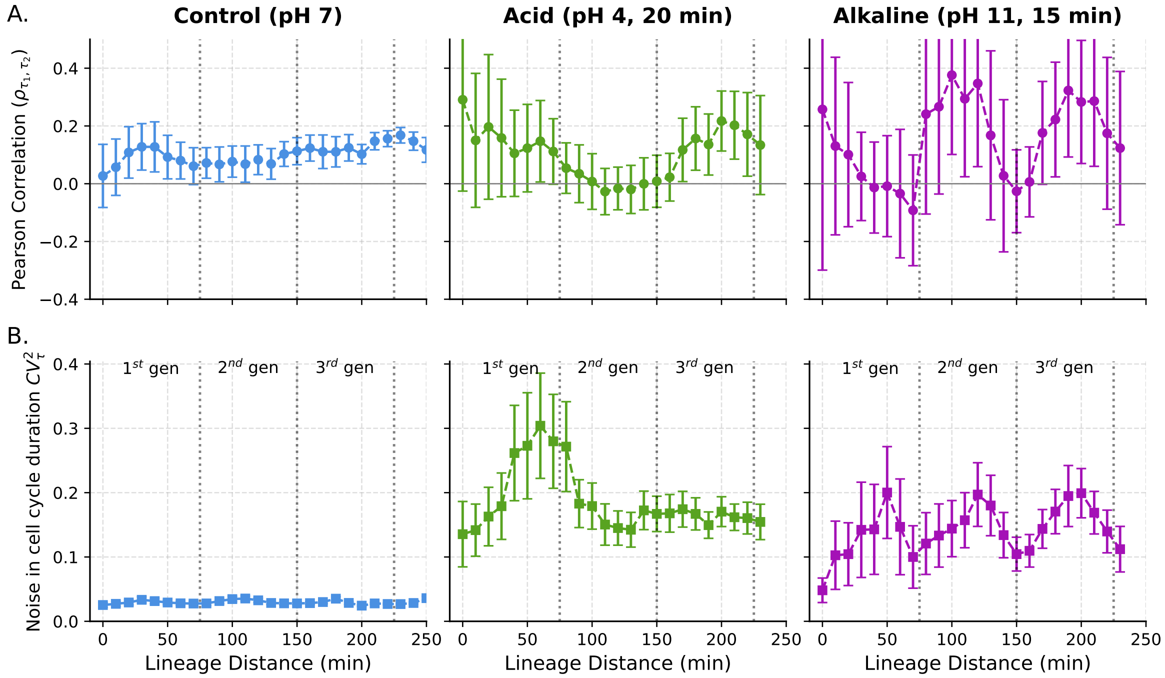


Figure S2: **Decomposition of the normalized mean squared difference into cycle-duration correlation and cycle-timing noise following transient pH stress.** Statistical metrics quantify the degree of alignment of cell cycles for cells with different degrees of relatedness. As a reference, we present cells under steady conditions (left: control, pH 7) and after transient environmental shocks (center: acid pulse, pH 4 for 20 min; right: alkaline pulse, pH 11 for 15 min). **(A)** Pearson correlation coefficient ($\rho_{\tau_1,\tau_2}$) of cell-cycle durations between lineage-related cell pairs as a function of lineage distance. The decay in correlation illustrates the gradual loss of transgenerational memory and division synchrony across the lineage tree. **(B)** Phenotypic noise of the population's cell-cycle duration, quantified by the squared coefficient of variation ($CV_\tau^2$). Data points represent a moving average computed with a $\pm$30-minute sliding window. Error bars indicate 95% confidence intervals derived from bootstrap methods. Vertical dashed lines mark the approximate duration of a typical cell cycle before stress ($\approx$ 75-minute intervals).

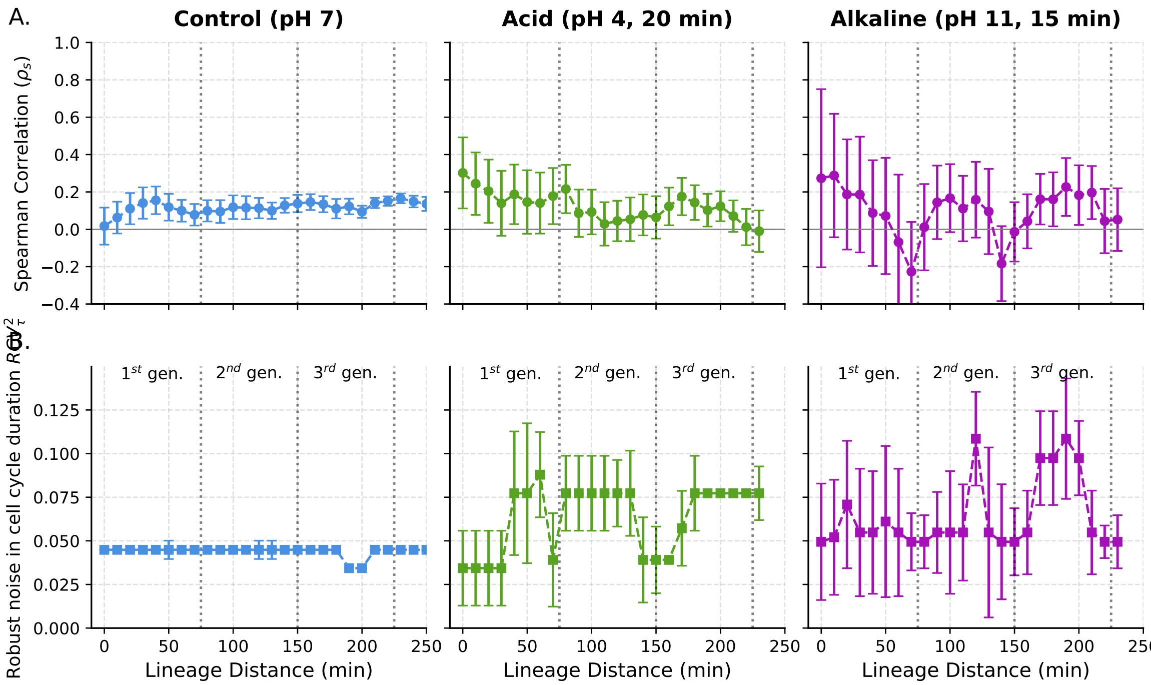


Figure S3: **Components of the cell-cycle difference computed with robust metrics of correlation and variability.** Statistical metrics quantify the degree of alignment of cell cycles for cells with different degrees of relatedness. As a reference, we present cells under steady conditions (left: control, pH 7) and after transient environmental shocks (center: acid pulse, pH 4 for 20 min; right: alkaline pulse, pH 11 for 15 min). **(A)** Spearman correlation coefficient ($\rho_{\tau_1,\tau_2}$) of cell-cycle durations between lineage-related cell pairs as a function of lineage distance. The decay in correlation illustrates the gradual loss of transgenerational memory and division synchrony across the lineage tree. **(B)** Phenotypic noise of the population's cell-cycle duration, quantified by the robust squared coefficient of variation ($rCV_\tau^2$). Data points represent a moving average computed with a $\pm$30-minute sliding window. Error bars indicate 95% confidence intervals derived from bootstrap methods. Vertical dashed lines mark the approximate duration of a typical cell cycle before stress ($\approx$ 75-minute intervals).
